# Assembly of hundreds of microbial genomes from the cow rumen reveals novel microbial species encoding enzymes with roles in carbohydrate metabolism

**DOI:** 10.1101/162578

**Authors:** Robert Stewart, Marc D. Auffret, Amanda Warr, Tim Snelling, Richard Dewhurst, Alan W. Walker, Rainer Roehe, Mick Watson

## Abstract

The cow rumen is a specialised organ adapted for the efficient breakdown of plant material into energy and nutrients, and it is the rumen microbiome that encodes the enzymes responsible. Many of these enzymes are of huge industrial interest. Despite this, rumen microbes are under-represented in the public databases. Here we present 220 high quality bacterial and archaeal genomes assembled directly from 768 gigabases of rumen metagenomic sequence data. Comparative analysis with current publicly available genomes reveals that the majority of these represent previously unsequenced strains and species of bacteria and archaea. The genomes contain over 13,000 proteins predicted to be involved in carbohydrate metabolism, over 90% of which do not have a good match in the public databases. Inclusion of the 220 genomes presented here improves metagenomic read classification by 2-3-fold, both in our data and in other publicly available rumen datasets. This release improves the coverage of rumen microbes in the public databases, and represents a hugely valuable resource for biomass-degrading enzyme discovery and studies of the rumen microbiome

## Introduction

New candidate enzymes for the efficient breakdown of plant matter are in high demand by biofuels and biotechnology companies, and the currently available hydrolytic enzymes are a potential limitation of our ability to produce bioethanol. The cow rumen is a fascinating organ, adapted for digesting cellulosic plant material and extracting energy and key nutrients. The rumen contains a microbial ecosystem in which a dense and complex mixture of protozoa, bacteria and anaerobic fungi convert carbohydrates to short-chain, volatile fatty acids (VFA).

Despite huge industrial and scientific interest, the rumen remains an under-characterised environment, containing many microbial species and strains that have not been cultured. In 2011, Hess *et al.*^1^ found that only 0.03% of their assembled metagenome had hits to sequenced organisms. Despite the fact that many thousands of sequenced bacterial genomes have been deposited in public repositories since then, metagenomic sequencing of the rumen still produces highly novel sequences which are significantly divergent from public collections^2,3^. These issues are significant barriers to identifying genes that encode the enzymes responsible for industrially relevant functions such as plant matter degradation.

Metagenomic binning^4^ is a cutting-edge bioinformatics technique that enables complete and near-complete bacterial genomes to be assembled directly from metagenomic sequencing data, without the need for culture, and has been applied to environments such as oil reservoirs^5^, aquifers^6^ and the human microbiome^7^. In addition, metagenome assembly and gene annotation tools have improved to enable bio-prospecting, the identification of novel proteins of industrial interest^8^.

Here we report the assembly of 220 complete and near-complete bacterial and archaeal genomes from a large rumen metagenomic sequencing study involving 42 Scottish cattle. We show that the addition of these genomes hugely improves our ability to quantify the taxonomic structure of the rumen microbiome, and many of the genomes encode novel carbohydrate-active enzymes that represent candidates of potential use in the biofuels and biotechnology industries.

## Results

### 220 draft microbial genomes assembled from the cow rumen

We produced 758 gigabases of Illumina sequencing data from 42 rumen microbiomes of Scottish cattle, and created a metagenomic co-assembly using MEGAHIT^9^. Contigs greater than 2 kb in length were used as input to metaBAT^10^ and CheckM^11^. The former clustered contigs based on tetranucleotide frequency and abundance across the 42-sample dataset; the latter assessed completeness and contamination of the resulting assemblies based on protein biomarkers throughout the taxonomic tree. Our analyses resulted in 220 high quality genomes with an estimated completeness greater than 80% and estimated contamination less than 10% (from now on referred to as RMGs, Rumen Meta-Genomes). Supplementary table 1 describes the assembly characteristics and predicted taxon of each genome, and Figure 1 shows a phylogenetic tree of the binned genomes alongside selected public genomes and the 15 binned genomes from Hess *et al.*^1^. Supplementary figure 1 shows a linear representations of the same tree. Supplementary table 2 shows the results of a comparison of the RMGs with public genomes using MinHash^12^ signatures, and supplementary table 3 provides summary results comparing the RMG proteomes with UniProt TrEMBL^13^. All of the above were used to predict the most likely taxon of each genome.

**Fig. 1.**
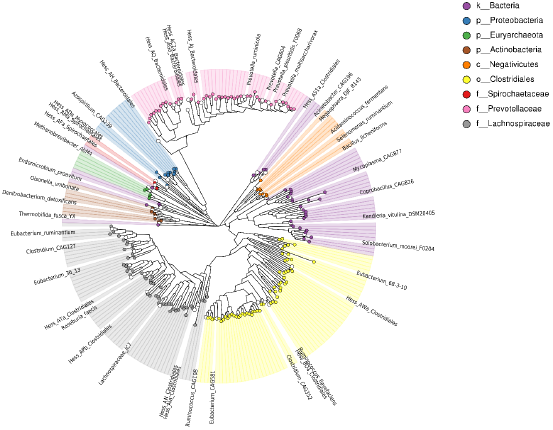
Phylogenetic tree of 220 draft genomes from the cow rumen, and closely related public genomes including 15 binned genomes from Hess *et al.* Coloured circles represent the RMGs. White circles represent public genomes and have correspond ding labels.

Several large clades are visible in figure 1. The tree is dominated by *Clostridiales*, with a large subset further identifiable as *Lachnospiraceae.* A large cluster of *Prevotellaceae* is also present. Smaller clades represent the *Proteobacteria, Archaea, Actinobacteria* and *Negativicutes*. The remaining nodes and branches represent miscellaneous bacteria.

We can confidently resolve five of the RMGs to species (>95% protein identity and near complete proteomes): RMG_87 is a strain of *Bacillus licheniformis*, RMG_788 a strain of *Kandleria vitulina*, RMG_720 a strain of *Acidaminococcus fermentans*, and RMG_623 a strain of *Megasphaera sp.* (most similar to strain *DJF_B143*). All four of these species have previously been found in ruminants^14-18^ and have traits of interest to the biotechnology, farming and breeding industries. *Bacillus licheniformis* is a species of huge interest to the biotechnology industry, encoding both hemicellulolytic and cellulolytic enzymes^14^ as well as serine proteases and other important enzymes. *Kandleria vitulina* was relatively recently renamed^19^, previously *Lactobacillus vitulinus* when isolated from bovine rumen in 1973^16^. *Kandleria vitulina* is a little-studied organism, though the family *Erysiopelotrichaceae* (of which it is a member) has been positively correlated with milk yield^20^ and negatively correlated with methane emissions^21^. *Acidaminococcus fermentans* is a Gram negative diplococcus that uses citrate as an energy source producing hydrogen and hydrogen sulphide^17^, which may explain its association with methane production in cattle^2^ (methanogens convert H_2_ to CH_4_). *A. fermentans* has also been shown to decrease the accumulation of tricarballylate, a toxic end product of ruminal fermentation, by oxidizing trans-aconitate^22^. *Megasphaera* spp. have been found in cattle, sheep and other ruminants. *Megasphaera elsdenii*’s ability to produce a variety of volatile fatty acids is of interest to the chemical industry^18^. Supplementation of the diet of dairy cows with *M. elsdenii*, which utilizes lactate as an energy source, has potential benefits for energy balance and animal productivity^23^. Finally, RMG_119 is a strain of *Thermobifida fusca*^24^, a likely contaminant from soil during grazing. We include it here for completeness and to improve the coverage of *Thermobifida* genomes in the public databases.

Of the remaining 215 RMGs, 16 were resolved to at least Genus, 105 to at least Family, 165 to Order, 172 to Class, 183 to Phylum and 215 to Kingdom. Five of the RMGs represent archaea, and supplementary figure 2 shows these in the context of 597 public archaeal genomes. We predict that all five RMGs represent methanogens due to their position in the tree. In each case, the closest matched organisms represent the most abundant and metabolically active methanogens in ruminants^25^. RMG_979 is most closely related to Candidatus *Methanomethylophilus*, a methanogen known to exist under anaerobic conditions^26^. RMG_576 and RMG_804 are related to *Methanosphaera* species. *Methanosphaera stadtmanae*, which RMG_576 is most closely related to, lacks many enzymes common to methanogens and relies on acetate for synthesis of cellular components. RMG_662 and RMG_565 are *Methanobrevibacter* species, also methanogens. *Methanobrevibacter boviskoreani*, which RMG_565 is closely related to, has previously been isolated from the rumen of Korean cattle^27^. The ratio of 5 archaeal genomes to 215 bacterial genomes is broadly in line with previous results calculating archaea:bacteria ratio in ruminants^2,28^. Of the RMGs resolvable to phylum level, *Firmicutes* dominated (69%), followed by *Bacteroidetes* (19%), *Proteobacteria* (3.6%), *Actinobacteria* (4.2%), *Euryarchaeota* (3%), *Spirochaetes* (0.6%) and *Tenericutes* (0.6%).

**Fig 2.**
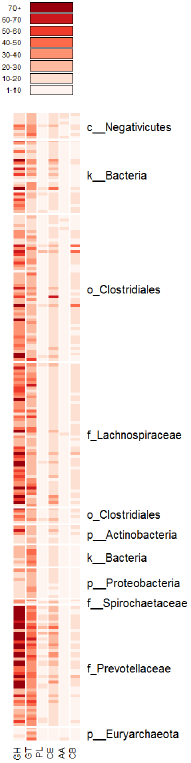
Distribution of carbohydrate-active enzyme classes across the 220 RMGs. GH=glycoside hydrolase, GT=glycosyl transferase, PL=polysaccharide lyases, CE=carbohydrate esterases, AA=auxiliary activities, CB=carbohydrate binding. White=absent, Dark Red=abundant.

### Thousands of novel carbohydrate-active enzymes

The carbohydrate-active enzymes database (CAZy^29^) defines six classes of enzyme involved in carbohydrate metabolism. Glycoside hydrolases hydrolyse the glycosidic bonds of complex carbohydrates and, within microbes, often assist in the degradation of cellulose, hemicellulose and starch; glycosyl transferases catalyse the formation of glycosidic bonds, utilizing sugar phosphates as donors, and transferring a glycosyl group to a nucleophilic group (together these two classes of enzyme form the major machinery for the breakage and synthesis of glycosidic bonds). Polysaccharide lyases cleave glycosidic bonds in polysaccharides; and carbohydrate esterases catalyze deacylation of polysaccharides. Finally, the Auxiliary Activities class within CAZy describes a number of enzymes that act in conjunction with the first four classes, and the Carbohydrate-Binding Modules (CBMs) describe proteins that adhere to carbohydrates.

The 220 RMGs contain 473,645 predicted proteins. These were searched against the CAZy database using dbCAN and filtered using suggested cut-offs^30^ (suppl. table 4). A total of 13,761 sequences were predicted to have at least one carbohydrate-active function. The proteins were compared to nr, env_nr, m5nr^31^, UniProt TrEMBL^13^ and the Hess *et al*^1^. gene predictions to check for novelty (suppl. Table 5), and against Pfam^32^ to detect other domains (suppl. Table 6). Of the 13,761 proteins, only 1123 (8.2%) had a highly similar match in any of the above databases (>= 95% identity), indicating that 12,638 of our predicted carbohydrate-active proteins can be considered novel.

In total, and including proteins with multiple domains, the RMGs contain 7298 glycoside hydrolases, 4530 glycosyl transferases, 185 polysaccharide lyases, 1688 carbohydrate esterases, 57 proteins with auxiliary activities, and 611 carbohydrate-binding proteins. The distribution of these enzymes across the 220 RMGs can be seen in figure 2 and supplementary figure 3. Glycoside hydrolases and glysosyl transferases are enriched in the *Prevotellaceae, Lachnospiraceae* and other *Clostridiales*, whilst being largely absent from the *Archaea, Proteobacteria* and *Negativicutes*.

**Figure 3.**
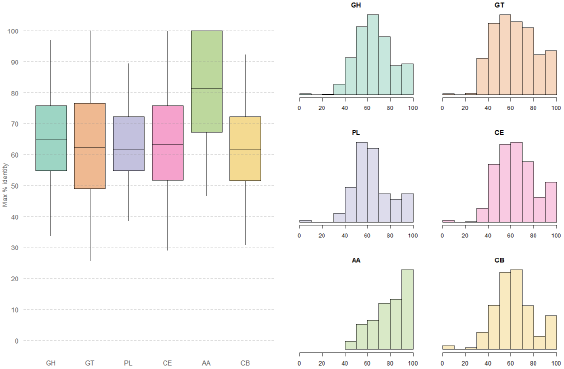
Distribution of the maximum percentage identity of the RMG proteins against five public databases for six classes of carbohydrate-active enzymes. GH=glycoside hydrolase, GT=glycosyl transferase, PL=polysaccharide lyases, CE=carbohydrate esterases, AA=auxiliary activities, CB=carbohydrate binding.

To understand how different the RMG proteins are from those in the public databases, we plotted the percentage amino-acid identity of the best hit for each CAZy enzyme class (fig 3). On average, the predicted glycoside hydrolases (GH), glycosyl transferases (GT), polysaccharide lyases (PL), carbohydrate esterases (CE) and carbohydrate-binding (CB) proteins are between 60 and 70% identical at the amino-acid level with current publicly available sequences. The auxiliary activities (AA) class are more conserved, with a median amino-acid identity just higher than 80%.

Finally, we investigated the ability of any of the RMGs to produce cellulosomes, multi-enzyme complexes responsible for the degradation of lignocellulosic biomass^33^. A key component of the cellulosome is a scaffoldin protein, which acts as a scaffold for the complex. The scaffoldin protein is characterised by multiple, repeated cohesin domains that the members of the cellusome bind to^34^. There are three RMGs that contains proteins with multiple predicted cohesion domains: RMG_290 has a protein with five predicted cohesion domains; RMG_556 a protein with 3; and RMG_334 a protein with two. According to our phylogenetic tree (fig 1 and suppl fig 1) RMG_290 is most similar to *Ruminicoccus flavifaciens*, one of the most dominant cellulolytic bacteria in the rumen^35-37^. However the average protein identity between RMG_290 and proteins in UniProt TrEMBL (which includes proteins from *Ruminicoccus flavifaciens)* is only 70.71%, indicating that RMG_290 is significantly different to existing *Ruminococcae* at the protein level. RMG_334 is classified as a member of the *Clostridiales*, and RIG_556 can only be classified as a member of the kingdom *Bacteria*, and is clustered closely with bin AS1a from Hess *et al.*^1^. All three genomes also contain glycoside hydrolases and glycosyl transferases.

### Improved metagenomic classification

Reads from this study, Hess *et al,.*^1^ and Shi *et al.*^38^ were taxonomically assigned to six different sequence databases using Kraken^39^ (see methods). The base database consisted of bacterial, archaeal, fungal and protozoan genomes from RefSeq^40^. Then each of GEBA^41^, and the RMGs were added, alone and in combination, and the effects on classification rate observed (Fig 4). In all three datasets, the base RefSeq database classifies fewer than 10% of the reads, and addition of the GEBA genomes has only a marginal effect. Addition of the RMGs increased the classification rate considerably. Overall classification rates were improved by 2.5- to 3- fold when adding rumen-specific bacteria from this study.

**Fig 4.**
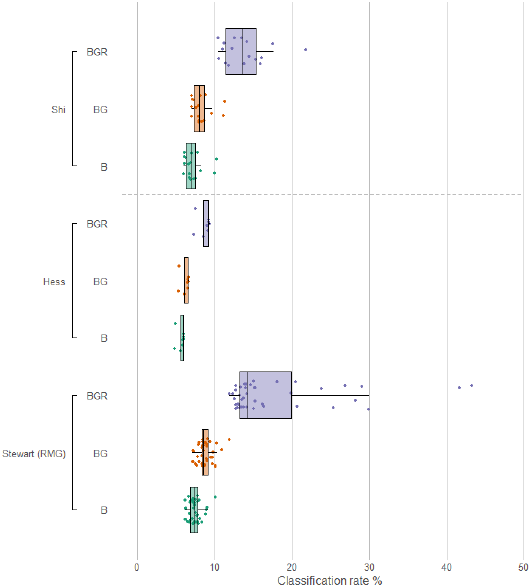
Classification rate for three datasets against various Kraken databases. B=bacterial, archaeal, fungal and protozoan genomes from RefSeq; G=1003 GEBA genomes; R=RMGs. Databases were used in combination. Addition of rumen-specific RMGs (R) has the most dramatic effect.

The classification rate for Hess *et al.* is noticeably lower than the other two datasets. We note that both our data and the Shi *et al.* data come from rumen fluid, whereas the Hess *et al.* data are specifically switch-grass associated, raising the possibility that these two microbiomes are significantly different. We re-classified all samples using Centrifuge^42^, an efficient metagenomic classifier capable of indexing the entirety of *nt*. Whilst the Kraken databases contain only microbial sequences, the *nt* database contains non-redundant sequences from all sequenced clades of life, and may be informative about the presence of non-microbial DNA in the Hess *et al.* samples. However, classification results against *nt* need be interpreted with care as many entries are mis-labelled or contain contamination. These analyses identified a number of factors which may explain the lower classification rates in the Hess *et al.* data. Firstly, the Hess *et al.* data have reads of length 100bp, whereas both our and the Shi *et al.* data are reads of length 150bp. The shorter read lengths may influence Kraken’s ability to find matches in the database. Secondly, as may be expected, the Hess *et al.* data included more reads originating from wheat and switchgrass (suppl. figure 4). Finally, *Prevotella* were generally more abundant in our data and the Shi *et al.* data, and *Fibrobacter* more abundant in the Hess *et al.* data (suppl figure 5). *Fibrobacter succinogenes* is a ruminant bacterial species that is known to associate with fibrous plant material^43^.

## Discussion

The RefSeq database now contains well over 75,000 prokaryotic genomes^44^, driven by decreasing sequencing costs^45^, improvements in bioinformatics techniques^4^ and large projects focused on specific environments^46,47^. However, the scale of microbial diversity is huge, and whilst continued deposition of thousands of bacterial genomes in public databases is welcome, their impact on studies of less sequenced environments is minimal. In this study we show that current RefSeq genomes are very poor at classifying reads from the rumen microbiome, and that the addition of 1003 microbial genomes^41^, specifically designed to broaden our knowledge of microbial life, has minimal effect. Only by sequencing microbes specifically from the rumen do we begin to see improvements in classification, with the genomes from this study improving rates 2.5- to 3- fold. The human microbiome project published thousands of novel microbial genomes, greatly increasing classification accuracy for metagenomics data from human-associated samples^46^. If other fields wish to match this, they must sequence similar numbers. As we show, metagenomic binning will be an important technique for improving our understanding of the genomic landscape of the rumen.

Eventually, if we are to fully understand the function of any microbiome, we must aim for 100% classification – in other words, to understand which genome each read comes from, and what functions those genomes encode. By assembling 220 genomes from our own dataset, and adding other rumen-specific genomes, some of our samples showed a classification rate above 40%. On average over 70% of reads were assembled in to contigs greater than 2 kb (see Methods), and therefore assigning taxonomies and functions to assemblies rather than the reads themselves could provide far greater insight. We have adopted a three-stage process in our microbiome research: stage 1 is discovery, simply sequencing what is there; stage 2 is association, correlating traits of interest with changes in the microbiome; and stage 3 is intervention, where we test microbial interventions to see whether we can alter those traits. From our work in this and previous studies^2^, it is clear that the rumen microbiome requires investment in discovery in order to maximise our potential to associate and intervene.

The rumen is of huge industrial interest due to its ability to release energy and nutrition from plant material. We show here that the 220 RMGs contain thousands of carbohydrate-active enzymes which differ significantly from existing representatives in the public domain. The RMGs contain over 12,000 novel proteins that are likely to be involved in carbohydrate metabolism, and which share on average only 60 to 70% amino-acid identity with similar protein sequences in the public domain. We further identify three RMGs that potentially encode the machinery to produce cellulosomes, multi-enzyme complexes that have high cellulolytic activity. Our ability to exploit these predictions is limited by our ability to characterise their function and activity at scale. Indeed, we believe it is one of the greatest challenges in biology and engineering – to design a system that can characterise the function of thousands of proteins simultaneously and cheaply. Such a system will require collaboration between biochemists, biologists, engineers, and synthetic and computational biologists, but could revolutionise fields as diverse as medicine, energy, manufacturing, biotechnology and conservation.

In this paper, we focus on bacterial and archaeal genomes, however fungi and protozoa are also key components of the rumen microbiome. Protozoa make up over 50% of the biomass of the rumen, yet their role remains unclear, as they are difficult to culture, study and sequence^48^. The important role of fungi in fiber degradation in the rumen has been known for decades^49^, yet again these species are under-studied and their genomes are not available. If we are to fully understand the rumen microbiome, these knowledge gaps need to be filled.

As we and others have shown, metagenomic binning is a powerful technique for the recovery of complete and near-complete microbial genomes without the need to culture. However, the assemblies remain fragmented, and it would be beneficial to completely assemble entire chromosomes. New sequencing technologies such as that offered by Pacific Biosystems and Oxford Nanopore^50^ are now able to produce long reads at reasonable scale and cost, and hybrid approaches enable complete chromosomal assemblies from complex bacterial genomes^51^. There is every reason to expect that hybrid short- and long-read sequencing will enable the complete end-to-end assembly of microbial chromosomes direct from metagenomic samples, and these approaches could revolutionise our understanding of complex microbiomes.

## Data availability

All raw data have been submitted to the European Nucleotide Archive (ENA) under project http://www.ebi.ac.uk/ena/data/view/PRJEB21624. Assembled genomes and proteomes are available from DOI: 10.7488/ds/2099

## Acknowledgements and funding

The Rowett Institute of Nutrition and Health and SRUC are funded by the Rural and Environment Science and Analytical Services Division (RESAS) of the Scottish Government. The Roslin Institute forms part of the Royal (Dick) School of Veterinary Studies, University of Edinburgh. This project was supported by the Biotechnology and Biological Sciences Research Council (BBSRC; BB/N016742/1, BB/N01720X/1), including institute strategic programme and national capability awards to The Roslin Institute (BBSRC: BB/P013732/1, BB/J004235/1, BB/J004243/1); and by the Scottish Government as part of the 2016-2021 commission.

Author statement

This pre-print has had certain analyses removed. This was at the request of data providers who have made their data public but have not yet formally published that data. We respect the wishes of people who make their data available ahead of publication.

**Supplementary table 1:** Description of the 220 Rumen Meta-Genomes (RMGs), including estimated taxon, CheckM completeness and contamination statistics, assembly size, N50, number of contigs and the longest contig length

**Supplementary table 2:** Summary data from comparison of the RMGs against various databases using MinHash signatures.

**Supplementary table 3:** Summary data from a comparison of RMG proteomes against UniProt TrEMBL. Each RMG has total number of predicted proteins, the number with hits, the number predicted to be full length (qlen / hlen > 0.8), the most popular genus (and the number of proteins hitting that genus), the most popular organism (and the number of proteins hitting that organism) and the average percentage identity across all hits

**Supplementary table 4:** Summary of the comparison of RMG proteomes against CAZy using dbCAN. Results show the relevant HMM hit, the length of the HMM, query ID and length, e-value, alignment details and the predicted coverage of the HMM in the protein

**Supplementary table 5:** Summary data from a comparison of the RMG CAZy hits against various public databases. hmm_hit is the predicted CAZy HMM. Proteins were searched against nr, env_nr, m5nr, and Hess *et al.* data. Values are the maximum percentage identity recorded.

**Supplementary table 6:** Full results from a comparison of RMG proteomes against Pfam using pfam_can

**Supplementary figure 1:** Linear representation of Figure 1 comparing the RMGs with themselves and public genomes. Created using PhyloPhlAn and FigTree

**Supplementary figure 2:** Comparison of five archaeal RMGs with 597 publicly available archaeal genomes. Created using PhyloPhlAn and FigTree

**Supplementary figure 3:** Heatmap showing distribution of predicted CAZy enzymes across the 220 RMGs

**Supplementary figure 4:** Predicted contamination from plants in our data and the Hess and Shi data, data from Centrifuge

**Supplementary figure 5:** Predicted distribution of *Prevotella* and *Fibrobacter* species in our data and the Hess and Shi data, data from Centrifuge

## Methods

### Samples

Animal experiments were conducted at the Beef and Sheep Research Centre of Scotland’s Rural College. The experiment was approved by the Animal Experiment Committee of SRUC and was conducted in accordance with the requirements of the UK Animals (Scientific Procedures) Act 1986.

The data were obtained from three cross breeds: Aberdeen Angus, Limousin, and Charolais and one pure breed: Luing. The animals were offered two complete diets *ad libitum* consisting (g/kg DM) of either 500 forage to 500 concentrate or 80 forage to 920 concentrate. As previously described in Roehe *et al*.^3^, the animals were slaughtered in a commercial abattoir where two post-mortem digesta samples (approximately 50 mL) were taken immediately after the rumen was opened to be drained. Illumina TruSeq libraries were prepared from genomic DNA and sequenced on an Illumina HiSeq 4000 by Edinburgh Genomics.

### Bioinformatics

Adapters were trimmed from the Illumina data using Trimmomatic^52^ and the subsequent trimmed reads used as input for MEGAHIT^9^. A 42-metagenome co-assembly was carried out using options --kmin-1pass, -m 60e+10, --k-list 27,37,47,57,67,77,87, --min-contig-len 300, -t16. The resulting assembly consisted of 22714724 contigs with an N50 of 1033 bp. Three filtered assemblies were created consisting of contigs >300 bp, >2000 bp and >5000 bp. BWA MEM^53^ was used to map reads back to the filtered assembly and Samtools^54^ used to calculate coverage. On average, 96% of reads mapped to the >300 bp assembly, 71% mapped to the >2000 bp assembly and 49% mapped to the >5000 bp assembly. The contigs greater than 2 kb in length were used as input to Metabat^10^, with options --minContigLength 2000, -- minContigDepth 2 and -B 20 (for ensemble binning). This produced 2683 bins, which were used as input to CheckM with options lineage_wf, -t 16, -x fa. After filtering for completeness >80% and contamination <10%, we were left with 220 near complete, high quality draft genomes. The predicted proteomes from each bin are produced by prodigal^55^ as part of the CheckM process. The proteomes of each bin were compared to UniProt TrEMBL^13^ using Diamond^56^, and the top hit, length and percentage identity recorded. This allowed us to predict the most likely genus for each contig within each bin. Genera not within the Bacteria or Archaea lineage were flagged as problematic (n=150), and contigs with these genera were removed from their respective bins using BioPerl^57^. The resulting “cleaned” bins were re-processed using CheckM, and the results can be seen in table 1. Ete3^58^ was used to expand the CheckM predicted taxon to a full taxonomy list. Genomes and gene predictions were loaded into a Meta4 database^59^.

The cleaned bins were compared, using MinHash sketches as implemented in Sourmash^60^, to 100,000 genomes in GenBank; and 1003 genomes from the GEBA project^41^. The results can be seen in supplementary table 1. The best hits from these analyses, combined with top hits from the UniProt searches above, were used to select publicly available genomes for comparison. Prodigal was used to predict proteins from these public genomes. A tree consisting of the 220 RMGs plus selected public genomes was calculated using PhyloPhlAn^61^ and visualised using GraPhlAn^62^ and Fig Tree^63^. This initial tree, which relates the RMGs to their most closely related public genomes, in combination with the outputs from CheckM, UniProt and Sourmash searches, allowed us to assign a taxon to each of the bins, and these can be seen in table 1. Trees were subsequently re-drawn with the updated taxonomic assignments and can be seen in figure 1.

Predicted proteins were compared to Pfam^32^ using pfam_scan.pl; the CAZy^29^ database using dbCAN^30^; and to nr, env_nr, md5nr^31^ and the Hess *et al.*^1^ predicted proteins using diamond^56^.

Reads were classified using Kraken^64^, against six custom databases: BFAP, consisting of 7318 complete bacterial genomes, 229 fungal genomes, 585 archaeal genomes and 75 protozoan genomes (all from RefSeq); BFAP+GEBA, consisting of the genomes from BFAP but with the additional 1003 genomes from the GEBA project^41^; BFAP+GEBA+RMGs, consisting of BFAP, GEBA and the 220 RMG binned genomes. Reads were also classified against the NT database using Centrifuge^65^.

